# Triosephosphate Isomerase as a Quantum Logic Gate: Could quantum decoherence be toxic?

**DOI:** 10.1101/2025.02.22.639452

**Authors:** Daniele Romanello, Andrea Romanello

**Affiliations:** Geriatrician, Internal Medicine, Ospedale San Pietro Fatebenefratelli, Rome; Via Castelfranco Veneto 33, 00191, Rome, Italy; Department of Information Engineering, Electronics, and Telecommunications Sapienza University of Rome Rome, Italy; Via Giulio Galli 41, 00123, Rome, Italy

**Keywords:** Quantum Biology, Triosephosphate Isomerase, Quantum Tunneling, Decoherence, Enzymatic Catalysis, Methylglyoxal, QM/MM Modeling, Metabolic Disorders, SGLT2 Inhibitors

## Abstract

This study presents the hypothesis that triosephosphate isomerase (TIM), a pivotal enzyme in glycolysis, functions as a quantum logic gate. Utilizing quantum mechanics, we model TIM’s catalytic conversion of dihydroxyacetone phosphate (DHAP) to glyceraldehyde-3-phosphate (G3P) as a quantum operation involving precise proton transfer. To explore the broader implications of this quantum behavior, we developed a quantum model to assess the impact of Sodium-glucose co-transporter 2 inhibitors (SGLT2i) on methylglyoxal formation, a toxic byproduct linked to advanced glycation end products (AGEs). Our model predicts that SGLT2i could reduce methylglyoxal by decreasing the likelihood of intermediate formation, providing a potential mechanism for their protective effects observed in clinical contexts, including vascular and renal protection in diabetes, nephropathy, and heart failure. By reframing TIM as a quantum logic gate, this study not only challenges traditional views of enzymatic function but also opens new avenues for quantum biology, offering profound implications for the future of metabolic disease research and drug development. Moreover considering methylglyoxal as a result of a quantum tunnel inefficiency, it’s possible to hypothesize a new “noxa patogena” explicating it’s action as quantum interference.

## Background

### Quantum Mechanics in Molecular Biology

Over the past decade, the intersection of quantum mechanics with molecular biology has yielded groundbreaking insights into the processes that govern life at the atomic level. Quantum biology, a rapidly emerging field, explores how phenomena such as superposition, tunneling, and entanglement (core principles of quantum mechanics) manifest in biological systems (Table 1). These effects, once considered relevant only in physics, are now recognized as playing crucial roles in a variety of biological functions.

**Table 1.**
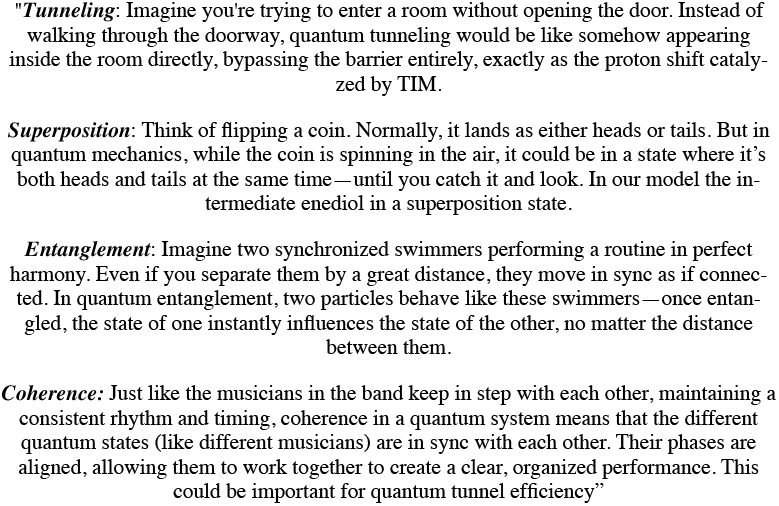

**Table 2.**
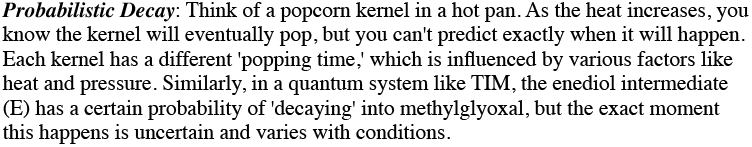

One of the milestone in quantum biology is the role of quantum coherence in photosynthesis. Research has demonstrated that quantum coherence may facilitate the efficient transfer of energy within the light-harvesting complexes of plants, algae, and bacteria. This phenomenon suggests that quantum states can enhance biological efficiency beyond the predictions of classical physics [1,2].

Quantum entanglement has also been proposed to play a role in the magnetic sense used by migratory birds. Entangled electron pairs in cryptochrome proteins within the birds’ eyes may help them sense the Earth’s magnetic field, aiding in navigation [3,4]. Additionally, quantum tunneling has been implicated in the sense of smell; the vibrational theory of olfaction suggests that electrons may tunnel between energy states in response to the vibrational frequencies of odorant molecules, enabling the detection of different smells [5]. Furthermore, quantum mechanics may be involved in genetic mutations through proton tunneling within DNA base pairs. This phenomenon could lead to tautomeric shifts, potentially causing point mutations during DNA replication, with significant implications for understanding mutagenesis and the evolution of genetic material [6].

### Quantum Mechanics and Enzymes

Enzyme catalysis represents another critical focus area in quantum biology, particularly the role of quantum tunneling. Quantum tunneling occurs when particles such as protons or electrons pass through energy barriers that would be insurmountable under classical mechanics. This process has been observed in enzymes, where it contributes to enhance catalytic efficiency [7,8]. The role of quantum tunneling in enzyme catalysis has become a focal point in quantum biology, particularly in the case of *triosephosphate isomerase* (TIM). Studies have shown that the tunneling of protons significantly contributes to the efficiency of proton transfer steps within TIM. Two critical proton transfer steps have been identified: the rate-limiting transfer of a proton from the substrate’s C-alpha carbon to Glu-165 and an intrasubstrate proton transfer involved in the isomerization of the enediolate intermediate. Computational studies using variational transition-state theory with semiclassical ground-state tunneling found that while tunneling plays a significant role, it accounts for a modest increase in reaction rate at room temperature—approximately tenfold—though the effect becomes more pronounced at lower temperatures. Furthermore, kinetic isotope effect studies have provided additional experimental validation of proton tunneling in TIM’s catalytic mechanism. These studies reveal that the primary and secondary hydrogens in dihydroxyacetone phosphate (DHAP) exhibit coupled motion during the proton transfer process, consistent with the presence of tunneling. The large secondary kinetic isotope effects observed in TIM-catalyzed enolization reactions, which are significantly greater than those seen in non-enzymatic reactions, underscore the unique catalytic efficiency of TIM facilitated by tunneling. Together, these insights highlight the critical role of tunneling in TIM’s function and suggest that disruptions in this quantum mechanical process, such as decoherence, could lead to the production of toxic byproducts like methylglyoxal [8-11].

### Bridging Quantum Mechanics and Metabolic Pathways

Quantum biology is rapidly bridging the gap between quantum mechanics and molecular biology, offering novel insights into the fundamental processes of life. Understanding the quantum mechanisms that regulate biochemical processes opens the door to a deterministic approach in studying metabolic pathways. This involves the creation of quantum models aimed at predicting cellular responses to specific variables with unprecedented precision.

### Study Objectives

In this context, our study aims to evaluate TIM as a potential quantum logic gate (an component of a quantum computer), positioned at the crossroads between glycolysis and lipid metabolism and the potential advance in biochemical modeling possibilities and medical implications. We also developed a quantum model with finality of investigate the possibilities that quantum decoherence could be implicated in methylglyoxal formation.

## Introduction

TIM is a pivotal enzyme in cellular energy metabolism, catalyzing the reversible interconversion of DHAP and glyceraldehyde-3-phosphate (G3P) within the glycolytic pathway. This enzymatic process is essential for maintaining the balance between the two triose phosphates produced by the cleavage of fructose-1,6-bisphosphate (F1,6BP) by aldolase, a step critical for the efficiency of glycolysis, lipid metabolism, and overall cellular energy production [12]. TIM is renowned for its exceptional efficiency, operating at a rate near the diffusion limit, with a catalytic constant (k_cat/K_m) among the highest recorded for enzymes [13] (Figure 3). Beyond its central role in glycolysis, recent research suggests that TIM may have additional functions in cellular metabolism and redox regulation, potentially interacting with alternative metabolic pathways such as the pentose phosphate pathway [14]. Furthermore, TIM has been implicated in other critical processes, including the regulation of glycolytic flux under stress conditions and involvement in signaling pathways associated with cellular apoptosis and aging [15].

The catalytic mechanism of TIM involves a precise proton transfer from the C1 hydroxyl group of DHAP to the C2 carbon of the enediol intermediate, leading to the formation of G3P (Figure 1). This proton transfer is tightly regulated by the enzyme’s active site, ensuring that the reaction proceeds with high efficiency and specificity [16,17]. The substrate’s proper orientation and the precise positioning of catalytic residues within TIM’s active site are crucial for its function, enabling near-perfect catalytic efficiency [17]. While the equilibrium of the reaction typically favors the production of DHAP, the rapid consumption of G3P by glyceraldehyde 3-phosphate dehydrogenase (GAPDH) during glycolysis shifts the metabolic flow from DHAP to G3P [17]. On the other side DHAP is reduced by glycerol-3-phosphate dehydrogenase (GPDH) to glycerol-3-phosphate (Gly3P), a precursor for lipid synthesis. This pathway, although slower and less prioritized compared to glycolysis, plays a significant role in anabolic processes like triglyceride and phospholipid synthesis [12]. (Figure 3)

**Figure 1:**
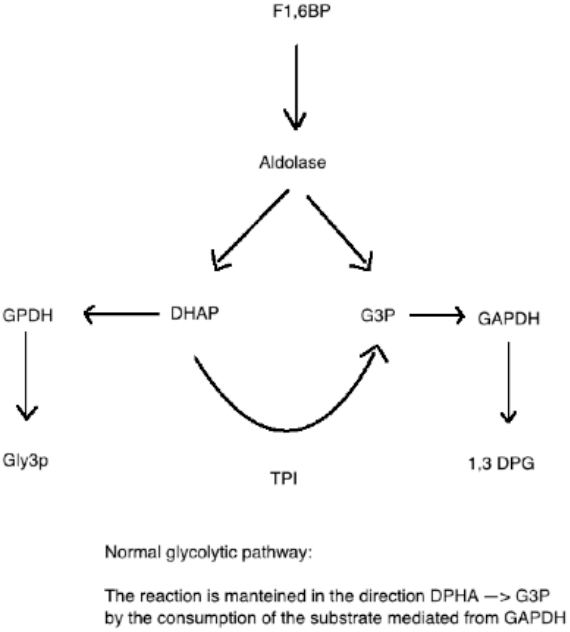

Recent advancements in quantum biology have opened new perspectives on how enzymes might operate at a quantum level. We assumed that TIM could operate as a quantum logic gate creating a parallelism between biological pathways and computing processes.

A quantum logic gate, a fundamental operation in quantum computing, manipulates qubits, the basic units of quantum information unlike classical logic gates, which operate on classical bits (0 or 1). A qubit can exist in a superposition of states, representing both 0 and 1 simultaneously [18]. This superposition allows quantum logic gates to perform complex computational operations that are impossible for classical gates, leveraging principles of quantum mechanics such as entanglement and superposition [18,19]. These operations are unitary, meaning they preserve the total probability and are reversible, a critical feature that allows quantum computations to be undone or reversed without loss of information [19,20].

For example, the Hadamard gate transforms a qubit into an equal superposition of 0 and 1, effectively enabling parallel computation. Similarly, the Controlled-NOT (CNOT) gate entangles two qubits, meaning the state of one qubit becomes dependent on the state of another [20]. This ability to entangle and superpose states underpins the power of quantum computing, enabling operations that exceed the capabilities of classical systems [19,20].

Given that, we propose that TIM could manage the equilibrium between molecular states in a manner similar to how a quantum logic gate manipulates qubits [20]. The precise proton transfers within TIM’s active site, akin to state transitions in a quantum system, might be governed by quantum mechanical principles, including superposition and tunneling [9,10].

Quantum tunneling, a phenomenon where a particle such as a proton can pass through an energy barrier that it classically shouldn’t be able to cross, could explain TIM’s extraordinary catalytic efficiency and the energy neutral reaction. In the context of TIM, this would allow the proton to move rapidly and efficiently between different positions within the molecule, facilitating the isomerization process [20]. The resulting conformational changes in the triosephosphate molecule are crucial because they determine the molecule’s metabolic pathway, thus regulating a key decision point in cellular metabolism [20].

Furthermore, the position of the proton after the reaction represents not just a physical change but also encodes biochemical information. This information, embedded in the structure of the product molecule, affects how it interacts with downstream enzymes [20]. Understanding TIM through this quantum lens challenges traditional views of enzymatic function and opens new possibilities for observing quantum effects in biological systems.

## Methods

### Quantum Mechanics - Molecular mechanics simulation (QM/MM)

We developed a quantum model (QM/MM) to conceptualizes TIM as functioning analogously to a quantum logic gate, influencing the equilibrium between DHAP, G3P and consequently, the levels of methylglyoxal. We also investigate the possibilities that decoherence can be involved in methylglyoxal production, a toxic byproduct of glucose metabolism.

### Computational Implementation

A Python script was employed to simulate the enzyme-catalyzed reaction, focusing on how variations in the number of iterations impact the probabilistic decay of the enediol intermediate. The script models the enzymatic process, specifically examining how decreased iterations influence methylglyoxal production.

#### Key Computational Steps

1. **Simulation of Proton Transfer**: The Python script simulates the transfer of a proton between DHAP and G3P, considering the quantum tunneling process that facilitates this transfer. The script calculates the likelihood of each proton transfer event, modeled as a quantum operation.
2. **Probabilistic Decay Analysis**: The enediol intermediate’s stability was analyzed through probabilistic decay functions. The script iteratively calculates the probability of the intermediate decaying into methylglyoxal, with the number of iterations.
3. **Impact of G3P levels reduction**: The model simulates the impact of SGLT2i on this process by altering the number of iterations and analyzing how this affects the probability of methylglyoxal formation. The reduction in G3P levels is simulated to understand the effect on the overall reaction dynamics.

### Quantum Logic Gate Framework

#### Quantum Logic Gates and TIM

TIM operate on qubits, manipulating their states according to the principles of quantum mechanics and governs the transition between molecular states (DHAP and G3P).

#### Properties of Quantum Logic Gates

- **Unitarity**: The operations are unitary, preserving the norm of the state vector and ensuring that the transition between states is reversible.
- **Reversibility**: The operations are reversible, allowing the system to return to its initial state without information loss.
- **Qubit Manipulation**: The gate operates on qubits, which can exist in superposition states, representing multiple potential outcomes simultaneously.

#### TIM’s Catalytic Function

- **State Transitions**: TIM catalyzes the interconversion between DHAP and G3P, with the enediol intermediate (E) representing a resonance state. The decay of this intermediate into methylglyoxal (M) represents a non-unitary, irreversible process.

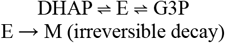

### Equilibrium Dynamics

The blockage of GAPDH reduces the metabolism of G3P, shifting the equilibrium towards DHAP formation. This increase in DHAP promotes the reverse reaction, potentially increasing the likelihood of methylglyoxal formation.

### Proton Positioning and Quantum Tunneling

In this model, TIM determines the proton’s position within the molecule. The positions DHAP and G3P represent two possible proton locations, while the enediol intermediate (E) represents a resonance state in which the proton is delocalized between the two positions. The conversion between DHAP and G3P via the intermediate E is conceptualized as a quantum tunneling event, wherein the proton passes through an energy barrier that would be insurmountable under classical mechanics.

### Validation and Future Directions

This model represents a theoretical framework that bridges quantum mechanics and enzymatic function. While the model provides compelling insights, experimental validation is essential. Future studies should focus on laboratory-based validation of the quantum behaviors hypothesized, particularly the role of quantum tunneling in TIM’s catalytic efficiency and its implications for methylglyoxal production. By simulating varying levels of TIM efficiency, this model could help guide experimental designs and enhance our understanding of enzyme function at the quantum level.

### Quantum Analysis

In this study, we treat TIM as a quantum logic gate, with its enzymatic function corresponding to transitions between quantum states. The key quantum states in this system are defined as follows:

- |**DHAP**⟩: The quantum state corresponding to dihydroxyacetone phosphate.
- |**G3P**⟩: The quantum state corresponding to glyceraldehyde-3-phosphate.
- |**E**⟩: The resonance intermediate state, representing a superposition of |DHAP⟩ and |G3P⟩.
- |**M**⟩: The quantum state representing the methylglyoxal, a toxic byproduct resulting from the decay of the enediol intermediate.

### Quantum States and Unitary Operators

The transitions between these quantum states are governed by unitary operators, ensuring that the quantum mechanical properties such as superposition and reversibility are preserved:

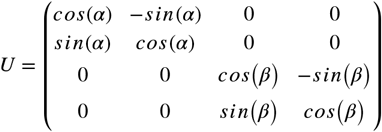

This unitary operator U describes the transitions between the states |DHAP⟩, |G3P⟩, and the resonance intermediate |E⟩. The angles α and β represent the transition probabilities, which can be adjusted based on the enzymatic environment and conditions.

### Decay Operator

The irreversible decay from the enediol intermediate |E⟩ to methylglyoxal |M⟩ is modeled using a Kraus operator, which accounts for the non-unitary, probabilistic nature of this process:

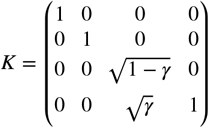

Here, γ represents the decay probability, which increases as the intermediate state |E⟩ persists. The Kraus operator K allows for the introduction of an irreversible pathway within an otherwise unitary framework, reflecting the real biochemical scenario where the enediol intermediate can either proceed to G3P or decay to methylglyoxal.

### Proton Position and Quantum Tunneling

The transition of the proton between |DHAP⟩ and |G3P⟩ via the resonance intermediate |E⟩ can be modeled as a quantum tunneling process. The efficiency of this tunneling is critical, as it minimizes the time the system spends in the intermediate state |E⟩, thereby reducing the likelihood of decay into the toxic byproduct methylglyoxal.

### Implementation Example

Consider the following scenario within our quantum model:

1. **Initial State**: Assume the system begins in the state |DHAP⟩.
2. **Unitary Evolution**: Apply the unitary operator U to simulate the transition to the resonance intermediate |E⟩ and further to |G3P⟩.
3. **Decay Process**: Introduce the Kraus operator K to model the probability of the system decaying from |E⟩ to |M⟩.

The evolution of the system can be described by the state vector ψ, which evolves as follows:

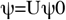

Where ψ0 is the initial state vector. After applying the unitary operator U for a specified number of iterations, the decay operator K is introduced probabilistically to simulate the formation of methylglyoxal.

### Final Considerations

This quantum analysis provides a novel framework for understanding the enzymatic action of TIM at a quantum level. By applying quantum mechanics to model the transitions between key molecular states, we gain insights into how TIM’s efficiency in catalyzing proton transfer impacts the production of toxic byproducts like methylglyoxal. This model not only advances our understanding of enzyme catalysis but also opens the door for future research into the quantum mechanical properties of biological systems.

### Dynamic Quantum State Evolution

We implemented a Python script to simulate the quantum state evolution of the system, focusing on the transitions between different molecular states (DHAP, G3P, and the enediol intermediate) and the probabilistic decay leading to methylglyoxal formation.

#### Biochemical Context

- **Aldolase Reaction:** The enzyme aldolase catalyzes the cleavage of F1,6BP into equal amounts of DHAP and G3P.
- **TIM Isomerization:** TIM exhibits a 20:1 preference for converting G3P into DHAP. This preference shifts the equilibrium towards DHAP under standard conditions.
- **GAPDH Activity:** GAPDH rapidly converts G3P into 1,3BPG during glycolysis, maintaining a dynamic equilibrium of approximately 98% DHAP and 2% G3P. This high activity level typically drives the reaction in the direction of DHAP to G3P.
- **GAPDH Blockage:** When GAPDH activity is inhibited, DHAP is metabolized by GPDH, albeit at a slower rate. This reduced velocity creates a backlog of triosephosphates, leading to an accumulation of DHAP and G3P.

#### Quantum State Simulation

The Python script developed for this study simulates the iterative processes involved in TIM-mediated isomerization. We modeled the effect of reactivating GAPDH and the subsequent reduction in G3P levels on the likelihood of isomerization and the formation of methylglyoxal. Specifically, the script analyzes the impact of varying the number of iterations (with a default of 100 iterations) on the probability of the enediol intermediate decaying into methylglyoxal.

### Python Script

The following Python script simulates the evolution of the quantum state ψ under the influence of unitary and Kraus operators, modeling the impact of reducing G3P levels on methylglyoxal production.

(Pyton Script is reported in appendices).

### Step-by-Step Explanation

1. **Operators Definition**:
  ○ **Unitary Matrix U**: This matrix represents the transitions between the states ∣DHAP⟩, ∣G3P⟩, and the enediol intermediate ∣E⟩. The angles α and β define the transition probabilities.
  ○ **Kraus Operator K**: This operator models the probabilistic decay from the enediol intermediate ∣E⟩ to methylglyoxal ∣M⟩. The decay probability is governed by γ, which represents the likelihood of the decay event occurring.
2. **Initial State**:
  ○ The system starts primarily in the state ∣G3P⟩, represented by the state vector `‘psi_0’`.
3. **State Evolution**:
  ○ The script runs a loop for `iterations = 100`, during which the state vector ψ is updated by applying the unitary transformation U to simulate the transitions between DHAP and G3P.
  ○ After each unitary transformation, the script calculates the probability Pd that the enediol intermediate will decay into methylglyoxal. This probability increases with each iteration.
  ○ If a decay event occurs (determined by a random number), the decay operator K is applied, altering the state vector ψ.
4. **Output**:
  ○ The script calculates and prints the probability of finding the system in the methylglyoxal state ∣M⟩ at the end of the simulation. This probability provides insights into how changes in TIM efficiency, represented by the unitary and decay operators, influence the formation of toxic byproducts

### Impact of Varying the Quantity of G3P on Methylglyoxal Formation

The quantum model with probabilistic decay suggests that the concentration of G3P plays a critical role in the formation of methylglyoxal (M). Specifically, reducing G3P levels leads to a corresponding decrease in the amount of the enediol intermediate (E), due to the relationship:

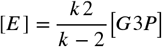

As a result, the production of methylglyoxal (M) diminishes, as indicated by:

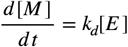

In this context, lowering G3P levels directly decreases the probability of E decaying into M. This relationship is preserved within the quantum model with probabilistic decay, as the decay probability Pd is contingent upon the concentration of E, which is in turn dependent on the availability of G3P.

The model illustrates that by reducing G3P, the system inherently reduces the likelihood of methylglyoxal formation, thus mitigating the production of this toxic byproduct. This insight aligns with the broader therapeutic implications of SGLT2i, which by modulating glycolytic flux, could potentially decrease cellular toxicity and contribute to better clinical outcomes in diseases associated with elevated methylglyoxal levels.

### Experimental Validation of the Quantum Model

To validate the quantum model proposed in this study, it is crucial to employ experimental techniques that can directly observe and quantify the dynamics of proton transfer and the formation of the resonance intermediate within TIM. The following approaches are recommended for a robust validation:

#### 1. Time-Resolved Spectroscopy

Time-resolved spectroscopy is a powerful technique that allows the observation of transient states in enzymatic reactions. By using ultrafast laser pulses, it is possible to capture the fleeting intermediate states of the enzyme-substrate complex.

- **Implementation**: In the context of TIM, time-resolved infrared (IR) spectroscopy or time-resolved UV-visible spectroscopy could be used to detect the vibrational or electronic transitions associated with the proton transfer. The spectral data obtained could be analyzed to determine the lifetimes of these intermediate states, providing direct evidence of the quantum tunneling process.
- **Expected Outcome**: The identification of distinct spectral signatures corresponding to the enediol intermediate would support the quantum model’s prediction of a superposition state. Additionally, observing changes in these signatures under different conditions could demonstrate how the quantum tunneling process is influenced by external factors.

#### 2. Single-Molecule FRET (Förster Resonance Energy Transfer)

Single-molecule FRET is a highly sensitive technique that can measure distances at the nanometer scale within individual enzyme molecules. This technique could be employed to monitor the conformational changes in TIM associated with proton transfer between DHAP and G3P.

- **Implementation**: By labeling TIM with donor and acceptor fluorophores at specific sites, it would be possible to track the distance changes during the catalytic cycle. As the enzyme undergoes conformational changes due to proton transfer, the FRET efficiency would change, providing real-time data on the dynamics of the process.
- **Expected Outcome**: The FRET data could reveal the timing and extent of conformational changes that correspond to the quantum tunneling of the proton. Comparing these observations with the predictions made by the quantum model would allow for direct validation of the model’s accuracy.

#### 3. Correlation with Enzymatic Kinetics Data

In addition to direct observation techniques, it is important to correlate the quantum model’s predictions with traditional enzymatic kinetics data. This approach involves comparing the predicted rates of proton transfer and methylglyoxal formation with experimentally determined kinetic parameters such as kcat, Km, and Vmax.

- **Implementation**: Kinetic assays could be conducted to measure the rate of TIM-catalyzed reactions under various conditions. These experimental rates can then be compared to the rates predicted by the quantum model, particularly focusing on the efficiency of proton tunneling and the probability of methylglyoxal formation.
- **Expected Outcome**: A strong correlation between the kinetic data and the model’s predictions would further corroborate the model’s validity. Discrepancies could indicate areas where the model needs refinement or where additional quantum mechanical effects might be at play.

## Discussion

This paper proposes the novel hypothesis that TIM acts as a quantum logic gate. TIM’s enzymatic activity involves manipulating the triosephosphate by altering the position of a proton, thereby modifying the information encoded in the molecule’s three-dimensional structure within its isomer. The transfer of the proton between C1 and C2 positions, given the enzyme’s high efficiency, can be linked to quantum tunneling, a process where particles pass through energy barriers that would be insurmountable under classical mechanics. This bidirectional and reversible process gives rise to a resonance intermediate, analogous to a quantum superposition state, where the system simultaneously exists in multiple states. The energy-neutral characteristic of TIM’s catalyzed isomerization reaction further supports the hypothesis that quantum tunneling could be involved. This model hypothesizes that the probability of forming methylglyoxal (which corresponds to the decay of the enediol intermediate, E) is inversely related to the efficiency of TIM in generating the proton tunneling process. Thus, the probability of methylglyoxal formation can be modeled as inversely proportional to the efficiency of TIM in facilitating proton tunneling or more precisely, the probability of methylglyoxal formation is assimilable to the probability that the proton tunneling fails. This model also suggests that, under standard efficiency conditions, the probability of methylglyoxal formation decreases when there is a reduced likelihood of triosephosphate isomerization. Given that, after the proper experimental validation, methylglyoxal could be found the first byproducts derived by a quantum inefficacy of an enzyme and it would be challenging to understand the nature behind potential noise or decoherence effects in biological systems that might impact on enzymatic quantum efficiency.The hypothesis that TIM functions as a quantum logic gate, manipulating qubits that achieve new positions through tunneling and passing through a superposition state (resonance intermediate), opens the door to an unprecedented deterministic approach in understanding enzymatic function. By considering biological effectors like TIM as quantum entities, we can create increasingly accurate theoretical models that could shift molecular, biochemical, and medical research towards a design-based framework. Expanding our knowledge in this direction, focusing research on identifying operators and circuits, with the aid of quantum physics where classical approaches fall short, could lead to the design of increasingly complex and realistic models. This approach might eventually enable us to decode the cellular “machine language,” recognizing its computational capabilities and allowing interactions with biological mechanisms at the deepest level.

### Medical Implications

The ability to generate quantum-based theoretical models, in tandem with the advancement of quantum supercomputers, offers an unprecedented level of precision in research. Identifying similar mechanisms at the cellular level allows for increasingly accurate predictions of therapeutically relevant changes. This precision can significantly accelerate the design of molecules, drug development, and the planning of clinical studies, especially with the support of artificial intelligence and machine learning. Such advancements would lead to a considerable reduction in both the time and cost required to develop new therapies. Furthermore, if validated, this model could redefine methylglyoxal as a product of enzymatic quantum inefficiency. This reclassification would introduce a novel category of ‘noxa patogena’, identifying agents that induce quantum decoherence within biological systems.

Although theoretical and still requiring validation, this quantum model holds significant potential for future research. The present model was inspired by observation of the clinical effect demostrated by SGLT2 inhibitors (SGLT2i) into diabetes, nephropathy and heart failure [21-28]. The model in appendices provides a mechanistic understanding of how SGLT2i treatments could reduce the production of methylglyoxal mitigating the formation of advanced glycation end-products (AGEs), which contribute to vascular and renal damage.

The quantum model implemented in Python provides a foundational theoretical approach, offering a basis for laboratory validation. We developed a model calculating the potential effect of SGLT2i on reducing methylglyoxal formation by decreasing the number of iterations in the catalytic cycle and, consequently, reducing the probability of the decay of the resonance intermediate (model in the appendices). Consequently, the reactivation of GAPDH by the effect of SGLT2i reduces the availability of substrates for TIM. In conclusion, SGLT2i may reduce the probability of probabilistic decay by decreasing the likelihood that a quantum tunneling event will occur for each molecule.

However, while this theoretical model presents a compelling hypothesis, it is crucial that these predictions be validated through rigorous experimental studies. Experimental validation is necessary to confirm the relationship between SGLT2i, TIM, and methylglyoxal formation, as well as to further explore the quantum mechanical aspects of enzymatic function. Only through empirical evidence can we substantiate the model’s predictions and fully understand the implications of SGLT2i on the biochemical pathways involved. We proposed advanced techniques such as time-resolved spectroscopy, single-molecule FRET and correlation with enzymatic kinetics data, highlighting the extreme complexity in experimentally validating this model. This technical difficulty, along with the lack of experimental data and preliminary studies, are the main limitations of the study.

## Conclusion

This model, although theoretical, demonstrate that TIM can be considered as a quantum logic gate that works changing the information delivered by the 3D structure of the isoforms of the triosephosphate.

In definition TIM appears to be intricately connected to quantum processes on multiple levels:

1. **Quantum Mechanics:** TIM may facilitates the quantum tunneling of a proton, a fundamental quantum phenomenon.
2. **Quantum Molecular Biology:** TIM may enables the reversible generation of two distinct molecular states, involving the formation of a resonance intermediate, which can be seen as a quantum superposition state.
3. **Quantum Computing:** Through the manipulation of a proton (acting as a qubit), TIM could generates distinct information encoded in the molecular structure.
4. **Quantum Medicine:** The failure of quantum tunneling within TIM may lead to the production methylglyoxal, highlighting its potential medical significance.

This novel perspective, if validated, could have significant implications across various levels of scientific research. It has the potential to enable the development of theoretical models with unprecedented precision, potentially accelerating advancements in biochemical and medical research while optimizing costs.

After the proper experimental validation, methylglyoxal could be the first byproducts derived by a quantum inefficacy of an enzyme and it would be challenging to understand the nature behind potential noise or decoherence effects in biological systems that might impact on enzymatic quantum efficiency.

Looking ahead, we strongly encourage the scientific community to explore this hypothesis further through experimental studies and interdisciplinary collaborations. Such efforts are crucial for validating the concept of enzymes functioning as quantum logic gates and could significantly advance our understanding and manipulation of complex biological systems. If confirmed, this discovery could represent a significant turning point in biotechnology and personalized medicine, offering new opportunities for innovations that transcend our current understanding of life sciences.

Moreover, future research could focus on identifying the specific experimental techniques, such as advanced spectroscopy or quantum computing simulations, that would best test the proposed quantum characteristics of TIM. By grounding this theoretical model in empirical data, we can move closer to realizing its potential impact on the fields of quantum biology and medical research.

## Appendices

### Application of Quantum Model to SGLT2 Inhibitors and Methylglyoxal

The catalytic action of TIM in proton transfer during the interconversion of G3P and DHAP presents a compelling analogy to the operation of a quantum logic gate in quantum computing [9,10]. Both systems rely on the precise and efficient control of fundamental particles, protons in the case of TIM, and qubits in quantum logic gates. The enediol intermediate formed during TIM’s catalytic cycle can be conceptualized as an intermediate quantum state, which must be meticulously managed to ensure the correct biochemical outcome (Table 1). Disruption of this conversion process can lead to the spontaneous decomposition of the enediol intermediate into a reactive alpha-ketoaldehyde, resulting in the loss of a phosphate group and the formation of methylglyoxal [12].

This side reaction is not catalyzed by any enzyme but occurs due to the intrinsic chemical instability of the enediol intermediate. The formation of methylglyoxal is particularly significant, as it is a highly reactive compound that can modify proteins and nucleic acids, leading to the formation of advanced glycation endproducts (AGEs) and contributing to cellular damage [12]. This process underscores the critical importance of TIM’s precision in controlling proton transfer, as any inefficiency can have deleterious biological consequences.

#### Oxidative Stress, PARP1 Activation, and SGLT2 Inhibitors

Glucose excess lead to an increased cellular oxidative stress. The activation of Poly(ADP-ribose) polymerase 1 (Parp1) in response to oxidative stress represents a critical cellular event with profound implications for metabolic pathways. Elevated levels of reactive oxygen species (ROS) trigger DNA damage, leading to the activation of PARP1. This enzyme, in turn, catalyzes the poly-ADP-ribosylation of target proteins, a process that significantly depletes intracellular NAD+, a cofactor indispensable for various metabolic reactions, including glycolysis [12,21].

GAPDH, a key enzyme in glycolysis, relies on NAD+ to facilitate the conversion G3P into 1,3-bisphosphoglycerate (1,3DPG) [12]. The depletion of NAD+ due to excessive PARP1 activation hampers GAPDH activity, leading to a metabolic bottleneck in glycolysis. This disruption causes the accumulation of G3P, which in turn shifts the equilibrium of TIM’s reaction towards the production of DHAP [21]. Despite the conversion in Gly3p, the activity of GPDH si slower of the activity of GAPDH, resulting in an accumulation of triosephosphates [21] (Figure 2).

**Figure 2:**
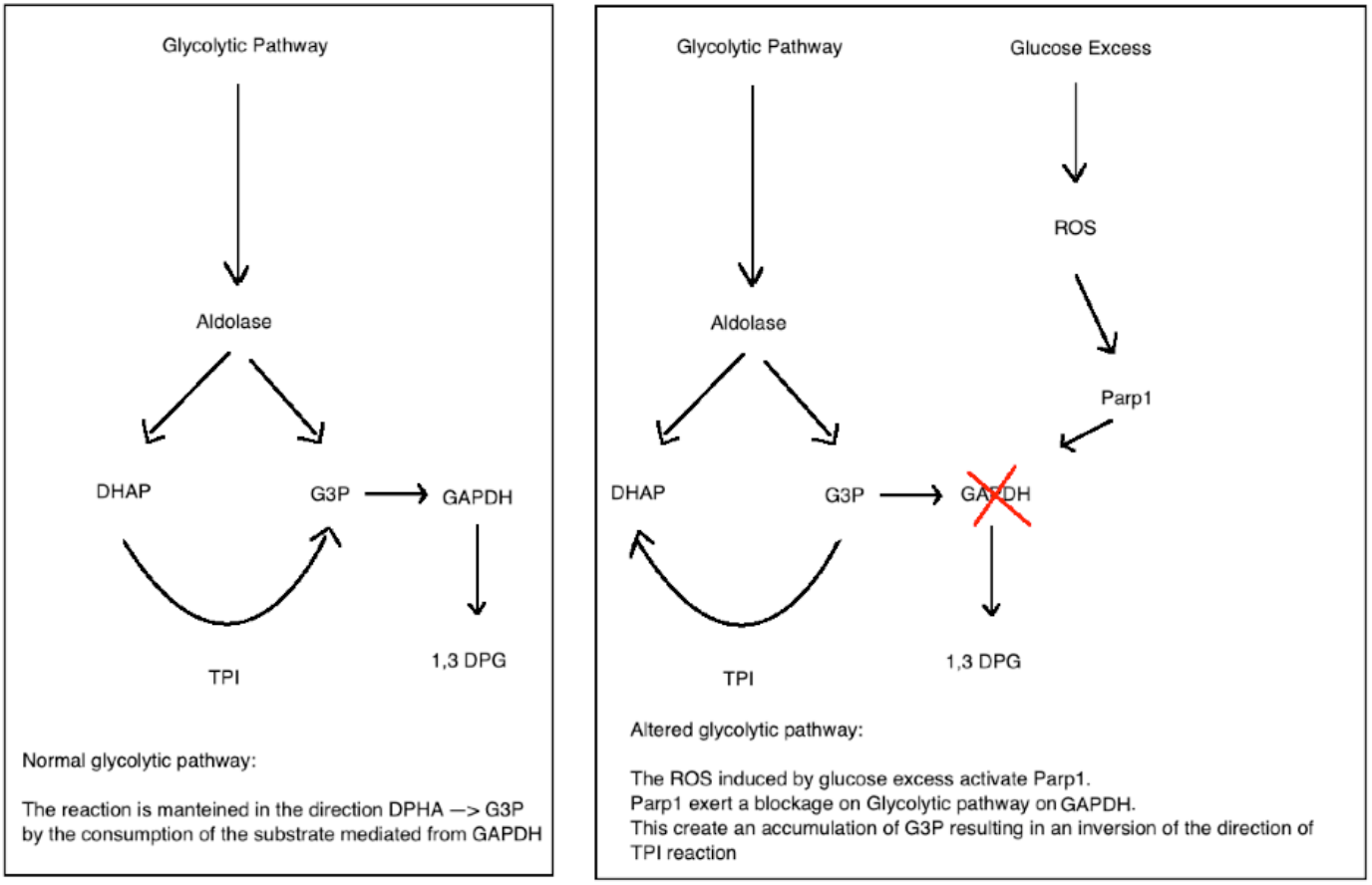

#### Impact of SGLT2 Inhibitors on Oxidative Stress and Glycolysis

SGLT2 inhibitors (SGLT2i), widely used in the treatment of type 2 diabetes and heart failure, are primarily known for their ability to lower blood glucose levels by inhibiting renal glucose reabsorption [22]. However, emerging evidence suggests that SGLT2i exert broader metabolic effects, particularly in modulating oxidative stress and attenuate the activation of PARP1, thereby preventing the depletion of NAD+ and restoring the functionality of GAPDH in glycolysis [21,26].

This restoration of GAPDH activity alleviates the metabolic bottleneck, decreasing the accumulation of G3P and thereby rebalancing the TIM-catalyzed interconversion between DHAP and G3P [12,21].

~~~
import numpy as np
# Definition of the operators
alpha = np.pi / 4 # Transition angle for U (unitary operation for DHAP to G3P)
beta = np.pi / 4 # Transition angle for U (unitary operation for G3P to DHAP)
gamma = 0.1 # Initial decay probability for K (probabilistic decay of the enediol intermediate)
# Unitary matrix U representing transitions between states
U = np.array([
  [np.cos(alpha), -np.sin(alpha), 0, 0],
  [np.sin(alpha), np.cos(alpha), 0, 0],
  [0, 0, np.cos(beta), -np.sin(beta)],
  [0, 0, np.sin(beta), np.cos(beta)]
])
# Kraus operator K representing the probabilistic decay of the enediol intermediate
K = np.array([
  [1, 0, 0, 0],
  [0, 1, 0, 0],
  [0, 0, np.sqrt(1 gamma), 0],
  [0, 0, np.sqrt(gamma), 1]
])
# Initial state vector psi_0, assuming the system is primarily in state B (|G3P⟩)
psi_0 = np.array([0, 1, 0, 0])
# Number of iterations to simulate temporal evolution
iterations = 100
# Initialize the state vecto
psi = psi_0
# State evolution simulation
for n in range(iterations):
  psi = U @ psi # Apply the unitary transformation U
  # Calculate the probability of decay
  P_d = 1 np.exp(-gamma * n)
  # Apply the decay operator K with probability P_d
  if np.random.rand() < P_d:
      psi = K @ psi
# Calculate the probability of finding the system in the methylglyoxal state (|M⟩)
prob_C = np.abs(psi[3])**2
# Output the probability of methylglyoxal formation
print(prob_C)
~~~

#### Declaration of generative AI and AI-assisted technologies in the writing process

During the preparation of this work the author used ChatGPT in order to improve readability. After using this tool, the author reviewed and edited the content as needed and take full responsibility for the content of the publication.

